# Broad-spectrum inhibition of *Phytophthora infestans* by root endophytes

**DOI:** 10.1101/107052

**Authors:** Sophie de Vries, Janina K. von Dahlen, Anika Schnake, Sarah Ginschel, Barbara Schulz, Laura E. Rose

## Abstract

- *Phytophthora infestans* (*Phy. infestans*) is a devastating pathogen of tomato and potato. It readily overcomes resistance genes and applied agrochemicals. Fungal endophytes provide a largely unexplored avenue of control against *Phy. infestans*. Not only do endophytes produce a wide array of bioactive metabolites, they may also directly compete with and defeat pathogens *in planta*.
- Twelve isolates of fungal endophytes from different plant species were tested *in vitro* for their production of metabolites with anti-*Phy. infestans* activity. Four well-performing isolates were evaluated for their ability to suppress nine isolates of *Phy. infestans* on agar medium and *in planta.*
- Two endophytes reliably inhibited all *Phy. infestans* isolates on agar medium, of which *Phoma eupatorii* isolate 8082 was the most promising. It nearly abolished infection by *Phy. infestans in planta*.
- Here we present a biocontrol agent, which can inhibit a broad-spectrum of *Phy. infestans* isolates. Such broadly acting inhibition is ideal, because it allows for effective control of genetically diverse pathogen isolates and may slow the adaptation of *Phy. infestans.*

## Introduction

*Phytophthora infestans* is a major pathogen of cultivated tomato (*Solanum lycopersicum*) and cultivated potato (*Solanum tuberosum*). Even today its impact cannot be ignored as it is still capable of destroying entire fields of its hosts, leading to up to 100% yield losses (Nowicki *et al.*, 2012). The two major control measures for *Phy. infestans* are resistance breeding and agrochemical applications. While several resistance genes have been identified in screens of wild relatives of *S. lycopersicum* and *S. tuberosum* (Song *et al.*, 2003; Van der Vossen *et al.*, 2003; Pel *et al.*, 2009; Zhang *et al.*, 2013), many of them are readily overcome by isolates of *Phy. infestans* (Vleeshouwers *et al.*, 2011). Similarly, agrochemicals can have a low durability in their protective function against *Phy. infestans* (Grünwald *et al.*, 2006; Childers *et al.*, 2015). Hence, continual scientific effort in terms of breeding, development of agrochemicals and other approaches, such as biological control, is needed for effective crop protection against this pathogen.

One approach that is gaining more and more attention is the use of endophytes for crop protection (Le Cocq *et al.*, 2016). Endophytes are microorganisms that grow within plants, and at the time of sampling, do not cause obvious symptoms on their host (Schulz & Boyle, 2005; Le Cocq *et al.*, 2016). Many studies have explored the bacterial, fungal and protist endophytic communities associated with different plants (e.g. Bulgarelli *et al.*, 2012; Lundberg *et al.*, 2012; Bodenhausen *et al.*, 2013; Schlaeppi *et al.*, 2013; Bulgarelli *et al.*, 2015; Edwards *et al.*, 2015; Busby *et al.*, 2016a; Coleman-Derr *et al.*, 2016; Ploch *et al.*, 2016). These studies indicate that the diversity of microbes living inside of plants is largely underestimated and that the distribution of some microorganisms is host and/or environment specific.

Furthermore, in some cases such endophytic microorganisms have been evaluated for their potential benefit to their hosts (Busby *et al.*, 2016b). Such benefits include growth promotion and protection against parasites and pathogens (e.g. Schulz, 2006; Lahlali & Hijri, 2010; Tellenbach & Sieber, 2012; Panke-Buisse *et al.*, 2015; Rolli *et al.*, 2015; Busby *et al.*, 2016a; Hiruma *et al.*, 2016; Martínez-Medina *et al.*, 2017). Often these functions are linked to metabolites produced and secreted by the endophytes (Son *et al.*, 2008; Puopolo *et al.*, 2014; Mousa *et al.*, 2016; Suryanarayanan *et al.*, 2016), highlighting the endophyte’s metabolic versatility(Schulz *et al.*, 2002; Strobel & Strobel, 2007; Verma *et al.*, 2009; Mousa & Raizada, 2013; Brader *et al.*, 2014). Endophytes may also directly compete with potential pathogens of their host plants (Alabouvette *et al.*, 2009), induce plant defense responses (Shoresh *et al.*, 2010) and/or produce bioactive anti-microbial metabolites (Brader *et al.*, 2014). An example of an endophyte that can be applied as a direct competitor of a plant pathogenic organism is *Phlebiopsis gigantea* (Adomas *et al.*, 2006). *Phl. gigantea* prohibits the infection of stumps of coniferous trees by the pathogen *Heterobasidion annosum sensu lato* and thereby limits the spread of the pathogen (e.g.Annesi *et al.*, 2005). Due to its success in limiting the spread of *H. annosum s.l.*, *Phl. gigantea* has been made commercially available. An example for the induction of defense responses by an endophyte is the barley root endophyte *Piriformospora indica*, which induces a jasmonic acid-dependent defense response in its host upon co-inoculation with a pathogen (Stein *et al.*, 2008). Furthermore, a recent study by Mousa *et al.* (2016) describes an *Enterobacter* sp. strain isolated from an ancient African crop (*Eleusine coracana* [finger millet]) with the ability to suppress the grass pathogen *Fusarium graminearum*. *Enterobacter* sp. traps *F. graminearum* in the root system of its host and simultaneously produces several antifungal compounds that kill the fungus.

Several bacterial and fungal endophytes, with the potential to inhibit *Phy. infestans* growth, have been described (Sturz *et al.*, 1999; Kim *et al.*, 2007; Miles *et al.*, 2012; Puopolo *et al.*, 2014). However, these endophytes have only been tested against single isolates of *Phy. infestans;* but alternative approaches, such as biocontrol, can show different outcomes depending on the pathogen isolate (Bahramisharif *et al.*, 2013). Therefore, the identification of endophytic species with a broad inhibition spectrum is of critical importance.

In this study, we screened the metabolite extracts of 12 fungal endophytes isolated from different plant hosts for their ability to inhibit growth of *Phy. infestans*. Using a plate assay with the four most successful fungal endophytes, we show that they inhibit the growth of a broad spectrum of European *Phy. infestans* isolates in co-culture. According to our phylogenetic analyses, these four endophytes are members of the Ascomycota. The endophyte with the strongest inhibition potential both on plates and *in planta* was *Pho. eupatorii,* isolate 8082. This endophyte prohibited proliferation of *Phy. infestans* and in some cases abolished its infection completely. Since we identified *Pho. eupatorii* based on the inhibition potential of its metabolite extract, the active component may be a secreted metabolite or a cocktail of different metabolites. A broad-spectrum activity as observed for *Pho. eupatorii* suggests either a conserved target for such secreted metabolite(s) or several targets that are specific for the pathogen isolate and that are covered by the complexity of the metabolite cocktail. Both can result in slower counter-adaptation of *Phy. infestans* to either the direct application of the endophyte or to the application of its metabolites. Therefore, *Pho. eupatorii* isolate 8082 is a potential novel broad-spectrum biocontrol agent of *Phy. infestans*.

## Material and Methods

### Isolation of endophytes

To isolate the endophytes, plant tissues of the respective hosts (Tab. S1) were first thoroughly washed under running water, then immersed for one minute in 70% ethanol, followed by 1-3min in 3% NaOCl and subsequently rinsed three times in sterile water. Sterilized tissues were imprinted on potato-carrot medium (Höller *et al.*, 2000) to test for effectiveness of sterilization and to optimize the sterilization procedure. The tissues were then cut with a sterile scalpel into 2mm slices and plated on potato-carrot agar medium with antibiotics (Höller *et al.*, 2000) and incubated for 3 weeks at 20°C. The emerging mycelia were taken into culture on potato-carrot agar medium and were initially identified according to morphology (Tab. S1).

### Screening crude metabolite extracts for anti-*Phytophthora infestans* activity

To test the growth inhibition potential of the 12 fungal endophytes, the endophytes were first grown on barley-spelt medium (Schulz *et al.*, 2011) and/or biomalt agar medium (Höller *et al.*, 2000) at room temperature for 21 days. To isolate the secondary metabolites, the cultures were extracted with ethyl acetate. 25μl of culture extract (40 mg/ml) were then applied to a filter disc and placed onto rye agar medium that had been inoculated with *Phy. infestans* isolate D2; subsequent incubation was at 20°C in the dark (Schulz *et al.*, 2011). Only fungal endophytes whose culture extracts resulted in a zone of inhibition ≥ 20mm were used for further analyses.

### Co-culture on plates

The fungal endophytic isolates 8082, 9907 and 9913, whose culture extracts had inhibited *Phy. infestans* in the agar diffusion assays and *Phialocephala fortinii* isolate 4197 (Schulz, 2006) were tested for their bioactivity against nine isolates of the late blight pathogen *Phy. infestans* (NL10001, NL88069, NL90128, IPO-C, IPO428-2, 3928A, D12-2, T15-2 and T20-2). The *Phi. fortinii* isolate was included based on previous experiments (Schulz *et al.*, 2002; Schulz, 2006; Schulz, unpublished). The co-culture experiments were performed and evaluated according to Peters *et al.* (1998). Fungal endophytes and *Phy. infestans* isolates were grown on rye-sucrose agar (RSA, Caten & Jinks, 1968) at room temperature. The duration of the experiments was dependent on the endophytes’ growth rates: eight days for all co-cultures that included 9913 and 14 to 16 days for the remaining co-cultures. A minimum of ten plates were analyzed per treatment. The Mann-Whitney U test (Mann & Whitney, 1974) was used to determine if differences between co-culture and control plates were significant. Average growth inhibition was estimated as: 1-(average radius in co-culture / average radius in control conditions). All experiments were evaluated again after eight weeks of incubation to assess long-term effects. Pictures were taken with an EOS 70D camera (Canon).

### Co-inoculation *in planta*

The surfaces of the *S. lycopersicum* seeds were sterilized using 70% ethanol for 3sec, followed by ~5% NaOCl for 30sec. The sterilized seeds were washed three times with sterile water for 3min. Seeds were incubated in the dark on 1.2% H_2_O-agar with a day-night temperature cycle of 18°C /15°C (16h / 8h). After three days, the seeds were transferred to a day-night cycle with 16h light (166 ± 17μmol quanta*m-2*s-1). Temperature conditions were the same as before. Nine to 11 days post sterilization (dps), the germinated seedlings were transferred to 9mm petri dishes containing 0.5% MS-medium (Murashige & Skoog, 1962) with 1% sucrose, poured as a slope.

An endophyte mycelial suspension was prepared from a two-to four-day old liquid culture for each endophyte (potato-carrot liquid medium; 100g potato-carrot mash [prepared according to Höller *et al.*, 2000] in 1l medium). Mycelium was equally dispersed in 25ml medium using Tissuelyser II (Qiagen, Hilden, Germany) for a few seconds. Preliminary inoculations of *S. lycopersicum* roots with 25 to 50μl of mycelial suspensions of all four endophytes were prepared. Endophyte isolate 9907 and *Phi. fortinii* killed the seedlings. Hence, only endophyte isolates 8082 and 9913 were used for further inoculation studies.

For inoculations with endophyte isolate 8082, 5μl or 10μl of the mycelial suspension or H_2_O (mock control) was applied to each root at 16dps. After 27dps seedlings were transferred to vessels (10cm×6.5cm×6.5cm) with MS agar medium. For inoculations with endophyte isolate 9913, 10μl of dispersed mycelium or H_2_O was applied to the roots of axenic seedlings at 18dps. However, the endophyte isolate 9913 did not grow sufficiently, so we performed a second inoculation with undispersed mycelium from the liquid culture at 22dps. These seedlings were transferred to vessels at 28dps. At 34 to 36dps each leaflet of endophyte and mock inoculated plants was inoculated with 10μl of *Phy. infestans* zoospore suspension (4°C cold) or with 10μl H_2_O (4°C cold). The zoospore suspension (5*10^4^ spores/ml) was harvested from a 25 days old culture of *Phy. infestans* isolate D12-2 and was kept on ice during the entire procedure. For the *Phy. infestans* zoospore isolation see de Vries *et al.* (2017). Plants were sampled for microscopic evaluation, to evaluate anthocyanin content and pathogen abundance at three days post inoculation with *Phy. infestans*.

To confirm endophytic colonization by the fungi, roots from the mock control, endophyte inoculated and co-inoculated samples were surface sterilized using three protocols: i) 70% EtOH for 3sec (isolate 8082) or 30sec (isolate 9913), ~5% NaOCl for 30sec, followed by washing three times with sterile H_2_O for 3min each (treatment 1), ii) 70% EtOH for 5min, 0.9% NaOCl for 20min, followed by washing three times with H_2_O (treatment 2, Cao *et al.*, 2004) and iii) 97% EtOH for 30sec, 10% NaOCl for 2min, followed rinsing four times with H_2_O (treatment 3, Terhonen *et al.*, 2016). Roots were imprinted on RSA agar plates to test for efficacy of sterilization and then placed on new RSA agar plates. The plates were evaluated at 8dps (isolate 8082) and 6dps (isolate 9913).

### Microscopy

Two aspects of host physiology were evaluated microscopically following the co-inoculation: chlorophyll intensity and relative necrotic area. Pictures to evaluate chlorophyll intensity were taken with the SMZ18 dissection microscope and a DS-Ri1 camera (Nikon, Tokyo, Japan) using a 600 LP filter (Transmission Filterset F26-010, AHF Analysetechnik, Tübingen, Germany), with an exposure time of 200ms and 100% gain. Intensity was measured using ImageJ2 (Schindelin *et al.*, 2015). Pictures for necrosis measurements were taken with a SteREO Discovery V8 binocular and an AxioCam ICc5 camera (Zeiss, Göttingen, Germany). The relative necrotic area was calculated as the necrotic area of a leaflet over the total area of the leaflet. The necrotic and total leaflet area were estimated using the ZEN Blue edition (Zeiss, Göttingen, Germany). Differences in relative necrotic area and chlorophyll content in the treatments were calculated using a Kruskal-Wallis test (Kruskal & Wallis, 1952) combined with a Tukey post-hoc test (Tukey, 1949) and using a Benjamini-Hochberg correction for multiple testing (Benjamini & Hochberg, 1995).

### Anthocyanin content evaluation

The anthocyanin content was measured and calculated according to Lindoo and Caldwell (1978). We analyzed three to six biological replicates per treatment. Samples were tested for normality using a Shapiro-Wilk test (Shapiro & Wilk, 1965) and for equal variance. Accordingly, significant differences were calculated using a two-sided t-test with the assumption of equal or unequal variances depending on the sample combination tested. All statistical analyses were done in R v 3.2.1.

### DNA and RNA extraction and cDNA synthesis

DNA was extracted from the mycelium of the fungal endophytes and *Phy. infestans* isolates grown on RSA medium using the DNeasy^®^ Plant Mini Kit (Qiagen, Hilden, Germany). RNA was extracted from infected and mock control leaflets of seedlings of *S. lycopersicum* using the Universal RNA/miRNA Purification Kit (Roboklon, Germany). Three to four leaflets were pooled per replicate. To evaluate RNA quality, 5μl of RNA were treated with 6μl deionized formamide, incubated at 65°C for 5min, followed by 5min incubation on ice. This mixture was then visualized on a 2% agarose gel. To ensure that no DNA contamination was present, the RNA was treated with DNAse I (Thermo Scientific). Reactions were adjusted for 200ng of total RNA. cDNA was synthesized with the RevertAid First Strand cDNA Synthesis Kit (Thermo Fisher Scientific, Lithuania).

### Molecular identification of endophytes

To determine the phylogenetic placement of the fungal endophytes, we sequenced their *internal transcribed spacer* (*ITS*) and 5.8S regions. *ITS1* and *ITS4* primers were used (White *et al.*, 1990). The 20μl PCR-reaction contained 1x Green GoTaq^®^ Flexi Buffer, 0.1mM dNTPs, 2mM MgCl_2_, 1U GoTaq^®^ Flexi DNA Polymerase (Promega, Madison, WI, USA), 0.2μM of each primer and 40-95ng of template DNA. The PCR protocol included an initial denaturation step of 95°C for 3min, followed by 35 cycles of a denaturation step at 95°C for 30sec, an annealing step at 60°C for 30sec and an elongation step at 72°C for 90sec, followed by a final elongation step of 72°C for 7min. All PCR products were purified with the peqGOLD Cycle-Pure Kit (Peqlab, Erlangen, Germany). The products were cloned into the pCR^™^ 4-TOPO^®^ vector of the TOPO^®^ TA Cloning^®^ Kit for Sequencing (Invitrogen, Carlsbad, CA, USA) and the plasmid DNA was extracted with the QIAprep Spin Miniprep Kit (Qiagen, Hilden, Germany). Sequencing was performed at Eurofins MWG Operon (Ebersberg, Germany). Sequences were blasted using BLASTn (Altschul *et al.*, 1990) and the best hits were retrieved. To assemble a dataset of closely related organisms from which to infer the phylogenetic placement of the unknown endophytes, the sequences of species with high similarity to our initial query sequences were downloaded. Taxonomic classification of these sequences was done using mycobank.org (provided by the CBS-KNAW Fungal Biodiversity Center, Utrecht). Additional sequences were retrieved from GenBank (Tab. S2). Taxonomically distant outgroups were chosen based on the systematic classifications in MycoBank (Crous *et al.*, 2004). The sequences were aligned using CLUSTAL-W and a Neighbor-Joining phylogeny was inferred using the Kimura-2 model with 5 gamma categories and pairwise deletion of gaps. One hundred bootstrap replicates were evaluated. All analyses were done using MEGA 5.2.2 (Tamura *et al.*, 2011).

### Assessment of endophyte and *Phytophthora infestans* growth after eight weeks of co-culture

To determine whether either the endophyte had overgrown *Phy. infestans* or *Phy. infestans* had overgrown the endophyte on the co-culture plates, we performed PCR reactions on DNA extracted from both sides of eight-week old co-cultures of five to nine *Phy. infestans* isolates with *Phi. fortinii,* isolate 8082 and isolate 9913 as well as their respective controls. We amplified the *ITS* loci (for primers see White *et al.*, 1990) and the *cytochrome oxidase subunit2* (*COX2*) using *Phytophthora*-specific primers from Hudspeth *et al.* (2000) with the protocol described above. Between 50-100ng of template DNA was used.

### Presence and abundance of *Phytophthora infestans*

To quantify the abundance of *Phy. infestans* in the seedlings pre-inoculated with the two endophytes (isolate 8082 and 9913) and the seedlings only inoculated with *Phy. infestans*, we performed a quantitative RT-PCR (qRT-PCR). The two markers, *PiH2a* and *PiElf1α*, were used for the pathogen and the three markers, *SAND*, *TIP* and *TIF3H,* were used as tomato (host) reference genes (de Vries *et al.*, 2015; de Vries *et al.*, 2017). Two independent qRT-PCR runs were used for the pathogen genes. All qRT-PCRs were performed in a CFX Connect^TM^ Real-Time System (Bio-Rad, Hercules, CA, USA) and included an initial denaturation at 95°C for 3min, followed by 40 cycles of a denaturation step at 95°C for 10sec and an annealing and elongation step of 60°C for 45sec. For *PiH2a* the annealing temperature was lower: 59°C in the first run and 55°C in the second run. For the following experiment, each run contained three biological replicates for: i) isolate 8082 (5μl mycelial suspension) with *Phy. infestans*, ii) isolate 9913 with *Phy. infestans* and iii) *Phy. infestans* without endophyte. Two biological replicates were completed for isolate 8082 (10μl mycelial suspension) with *Phy. infestans*. In each run, we analyzed three technical replicates for each biological replicate, resulting in six technical replicates for each biological replicate for both marker genes. To calculate the relative abundance of *Phy. infestans* in these samples, we set the Cq-values of those biological replicates that gave no biomass marker amplicon to 41. As the two independent runs gave the same results, they were combined. *PiH2a* and *PiElf1α* expression was then calculated according to Pfaffl (2001). Data were tested for normal distribution using a Shapiro-Wilk test and the appropriate statistical tests were then applied. For co-inoculations with isolate 8082, significant differences were calculated using a Mann-Whitney U-test. For co-inoculations with isolate 9913, significant differences were calculated using a two-tailed t-test. The statistical analyses were done using R v. 3.2.1.

## Results

### Metabolite screening identifies three endophytes with biocontrol potential

To identify fungal endophytes that, on the basis of their secreted metabolites, could be used as biocontrol agents against *Phy. infestans,* we screened culture extracts of 12 fungal endophytes for growth inhibition of *Phy. infestans* isolate D2 using an agar diffusion assay. Inhibition of *Phy. infestans* varied considerably, depending both on the endophyte isolate and on the culture medium. The average growth inhibition was 12.4 ± 8.7mm ranging from 0 and 35mm from the point of extract application (Tab. S3). Culture extracts of three of the 12 isolates inhibited growth of *Phy. infestans* with a radius ≥20mm. These three fungal endophytes (isolates 8082, 9907 and 9913) with the greatest *Phy. infestans* growth inhibition were chosen for further studies. An additional fungal strain, *Phi. fortinii* (isolate 4197) was included due to its mutualistic interaction with another host, *Larix decidua* (Schulz *et al.*, 2002; Schulz, 2006), growth inhibition of other pathogenic microbes and prior information that it could colonize *S. lycopersicum* asymptomatically (Schulz, unpublished).

### Phylogenetic placement of fungal endophytes

To determine the taxonomic identity and phylogenetic placement of the four selected fungal endophytes, we sequenced their *ITS1,* 5.8S and *ITS2* regions. First, we used these sequences in a BLAST search to identify the closest relatives of the fungal endophytes (Tab. S4). All four endophytes belong to the ascomycetes. Our analyses further supported the previous characterization of isolate 4197 as *Phi. fortinii* (99% identity and e-value 0, Grünig *et al.*, 2008). For isolate 8082 the best BLAST hit with 100% identity and an e-value of 0 was *Phoma eupatorii.* This was additionally supported by the fact that isolate 8082 was isolated from *Eupatorium cannabinum* (Tab. S1). The placement of isolates 4197 and 8082 in our phylogenetic analyses together with the extremely short branch lengths to their best BLAST hits further support these phylogenetic assignments (Fig. 1a,b). The best hit for isolate 9907 was *Pyrenochaeta cava* (95% identity and e-value 0) and for isolate 9913 it was *Monosporascus ibericus* (97% identity and e-value 0). This suggests that no completely identical sequence/taxa are currently represented in the database. *Pyrenochaeta* does not form a monophyletic group within the order of Pleosporales (Zhang *et al.*, 2009; Aveskamp *et al.*, 2010; Fig. 1c), thus based on the phylogenetic analyses isolate 9907 can only be placed within the order Pleosporales. Isolate 9913 was isolated from the roots of *Aster tripolium*, a plant that was growing in the salt marshes of the Mediterranean Sea (Tab. S1). Of note is that *Monosporascus ibericus,* the fungal endophyte clustering most closely with isolate 9913 in the phylogenetic analysis, has been recently described as an endophyte of plants growing in environments with high salinity (Collado *et al.*, 2002). Furthermore, the genus *Monosporascus* is monophyletic; isolate 9913 has been placed within this monophyletic group and herewith termed *Monosporascus* sp. (Fig. 1d).

**Figure 1.**
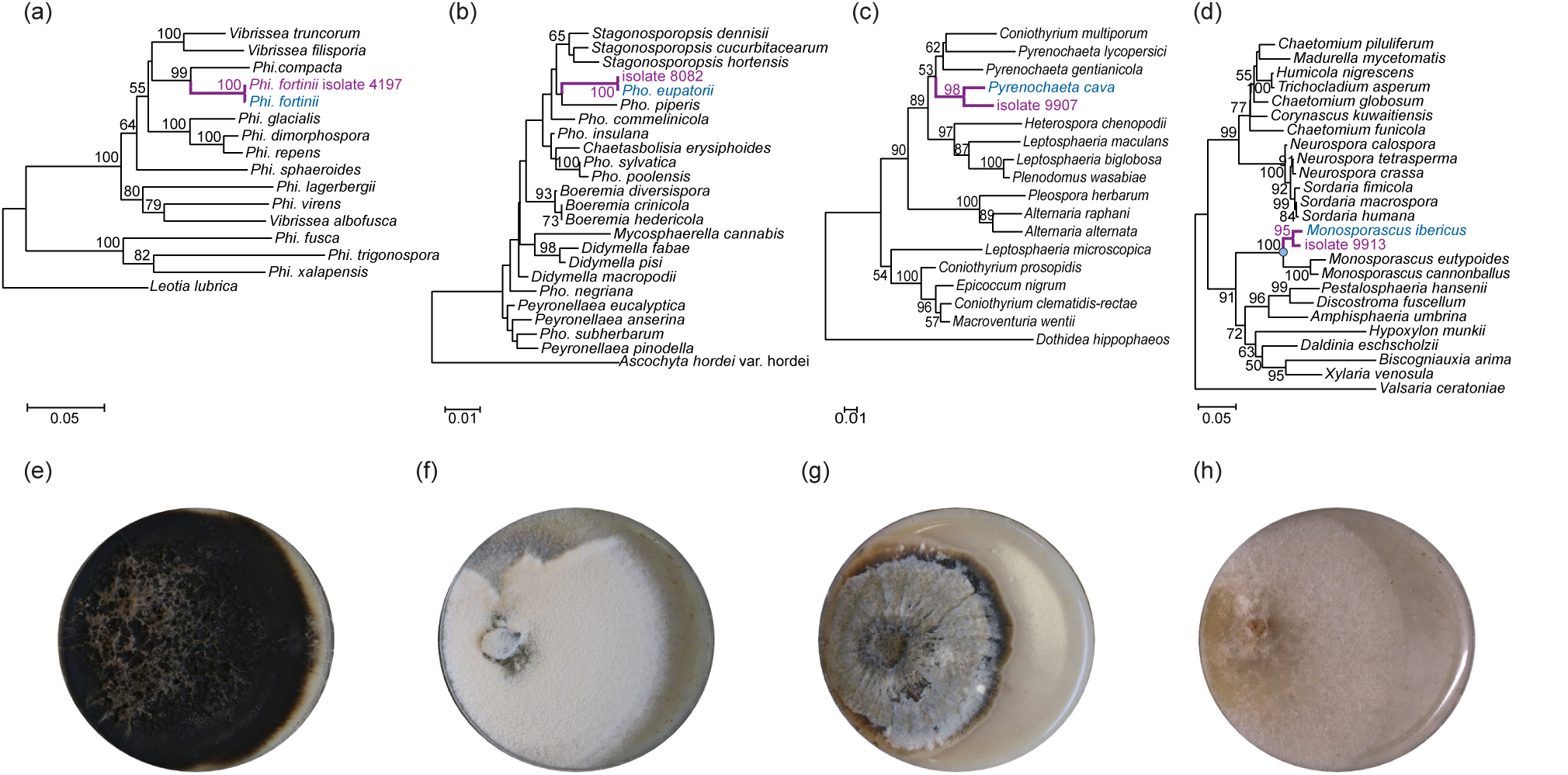
Phylogenetic placement of fungal endophytes. Neighbor-Joining phylogeny of ascomycetes closely related to the four fungal endophytes (a-d). Cloned sequences are shown in purple and the best BLAST hit is shown in blue. The monophyletic clade of the genus *Monosporascus* is indicated by the blue dot (d). The trees are rooted with *Leotia lubrica* (a), *Ascochyta hordei* var. hordei (b), *Dothidea hippophaeos* (c) and *Valsaria ceratoniae* (d). Only bootstrap values >50 are shown. The bar below the phylogeny indicates the distance measure for the branches. The corresponding fungal endophyte in culture is shown below each tree: *Phialocephala fortinii* (e), *Phoma eupatorii* (f), isolate 9907 (g) and *Monosporascus* sp. (h).

### Fungal endophytes show broad-spectrum inhibition of *Phytophthora infestans* growth

Our initial screening of the culture extracts identified endophytes with the potential to inhibit the growth of a single *Phy. infestans* isolate. We therefore wondered whether the inhibition could be effective against a wider range of isolates of *Phy. infestans.* To test this, we conducted a co-culture assay on RSA agar medium with the four fungal endophytes against nine European *Phy. infestans* isolates (Fig. 2). In the plate assay all four endophytes were capable of significantly restricting growth of *Phy. infestans* (Fig. 3). *Pho. eupatorii* and isolate 9907 showed a global inhibition of all *Phy. infestans* isolates tested (Fig. 3b,c). *Phi. fortinii* inhibited the growth of eight out of nine isolates and *Monosporascus* sp. inhibited the growth of seven of the nine isolates (Fig. 3a,d). *Pho. eupatorii* had the greatest average relative growth inhibition of *Phy. infestans* with 50.6 ± 2.2%, and *Monosporascus* sp. had the lowest with 11.9 ± 1.6% (Tab. 1).

**Figure 2.**
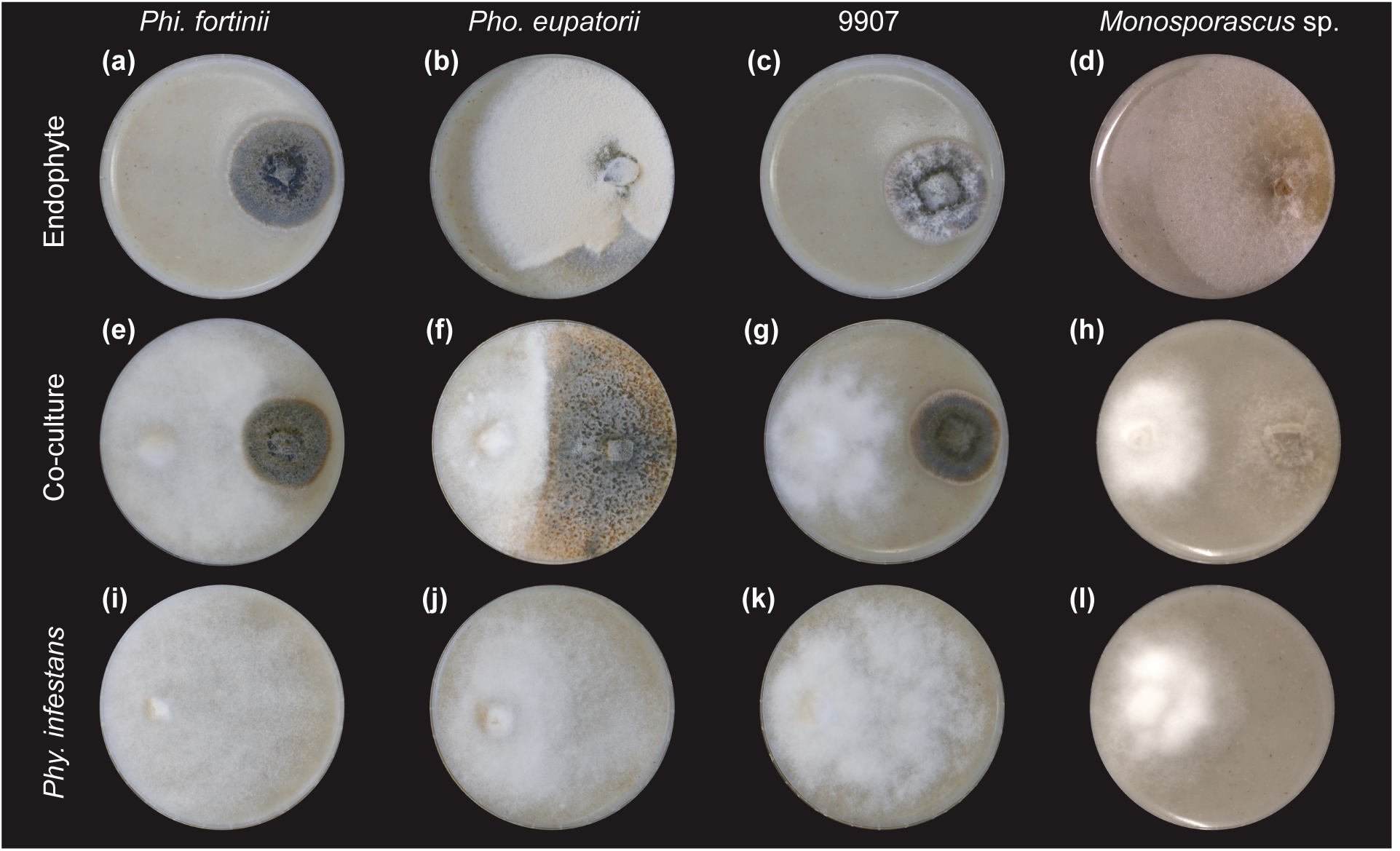
Co-cultivation of fungal endophytes with *Phytophthora infestans* on plate. Examples of two-week-old single and co-cultivations of *Phialocephala fortinii* with *Phy. infestans* isolate 3928A (a, e, i), *Phoma eupatorii* with *Phy. infestans* isolate NL90128 (b, f, j) and 9907 with *Phy. infestans* isolate T15-2 (c, g, k) and eight-day old single and co-cultivations of *Monosporascus* sp. with *Phy. infestans* isolate D12-2 (d, h, l). The diameter of each plate is nine cm.

**Figure 3.**
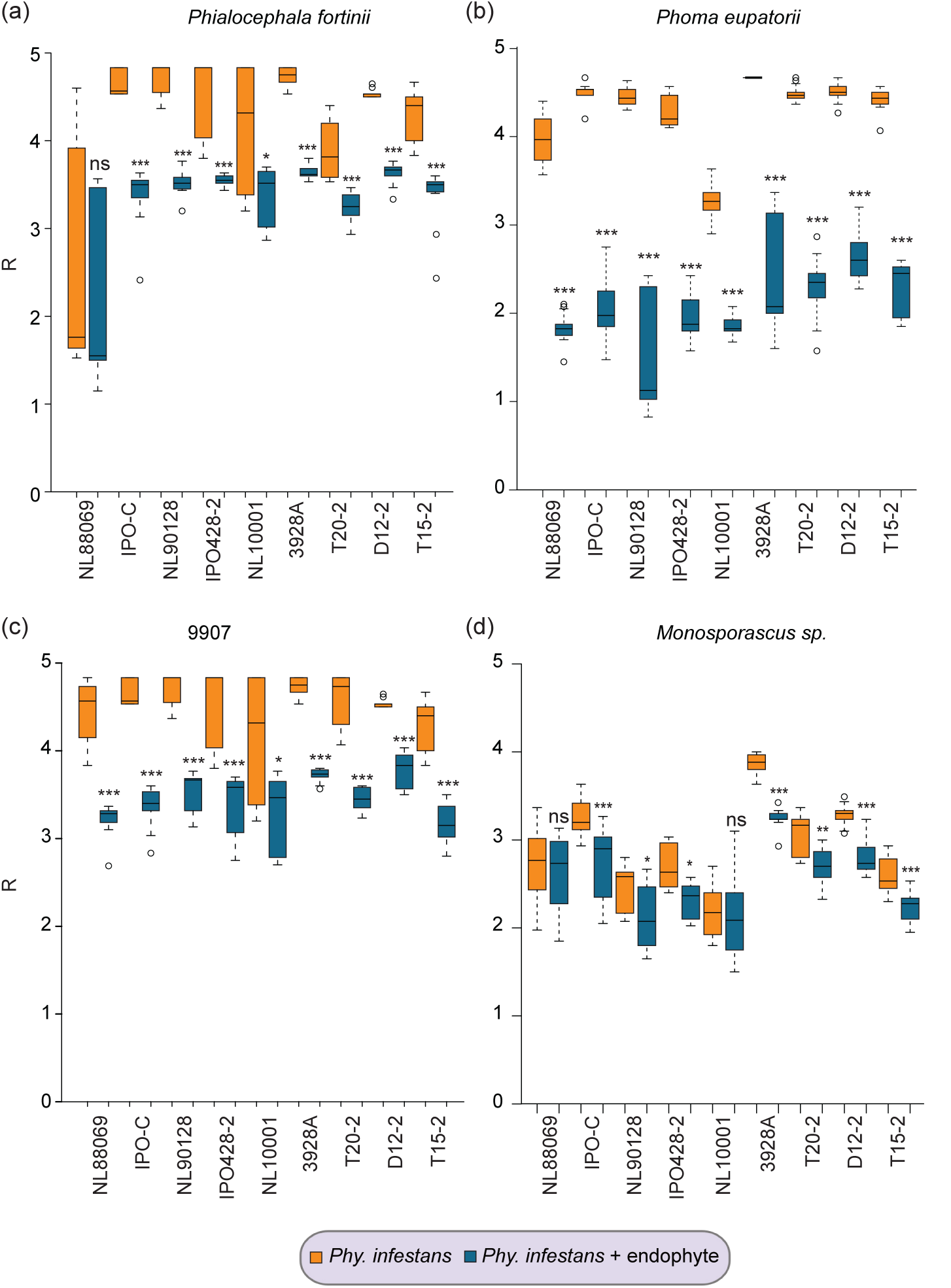
Radial growth inhibition of *Phytophthora infestans* isolates by fungal endophtyes. Radial growth (R) of the different *Phy. infestans* isolates denoted on the x-axis when grown alone (orange) or in dual culture with the four fungal endophytes (blue): *Phi. fortinii* (a), *Pho. eupatorii* (b), isolate 9907 (c) and *Monosporascus* sp. (d). The box indicates the upper and lower 50% quartile (interquartile range, IQR), the horizontal line in each box shows the median, the whiskers indicate the upper and lower bounds of the 1.5x IQR and the circles show data points, which are outliers. Significant differences are noted as * p< 0.05, ** p< 0.01, *** p< 0.001 and ns = not significant.

**Table 1.** Average relative growth inhibition of endophytes (first column) by *Phy. infestans* (upper row). The relative inhibition is calculated from the average radii estimated for co-cultivations and control plates.

|  | NL88069 | IPO-C | NL90128 | IPO428-2 | NL10001 | 3928A | T20-2 | D12-2 | T15-2 | Average+/- SEM |
| --- | --- | --- | --- | --- | --- | --- | --- | --- | --- | --- |
| *Phi. fortinii* | 0.060 | 0.109 | 0.119 | 0.139 | 0.143 | 0.145 | 0.016 | 0.188 | 0.077 | 0.111<br>+/-0.016 |
| *Pho. eupatorii* | 0.002 | 0.050 | 0.004 | 0.070 | 0.089 | 0.068 | 0.059 | 0.049 | 0.034 | 0.047<br>+/-0.009 |
| 9907 | 0.010 | -0.024 | 0.016 | -0.037 | 0.045 | 0.048 | 0.028 | 0.033 | 0.020 | 0.015<br>+/-0.009 |
| *Monosporascus*<br>sp. | 0.066 | 0.116 | 0.079 | -0.032 | -0.026 | 0.015 | 0.033 | 0.110 | 0.052 | 0.046<br>+/-0.017 |

To exclude a mere reduction based on growth limitations we i) measured the inhibition of the endophyte’s growth by *Phy. infestans* after the initial co-cultivation phase and ii) evaluated long-term co-cultures (i.e. eight weeks) to analyze the endophyte and pathogen growth progression. The growth of isolate 9907 was not inhibited by any of the *Phy. infestans* isolates (Fig. S1c). However, some isolates of *Phy. infestans* were able to inhibit the growth of the other three fungal endophytes (Fig. S1a,b,d). In all cases, the average relative inhibition of an endophyte by *Phy. infestans* was, however, less than the average relative inhibition of *Phy. infestans* by an endophyte (Tab. S5). For example, whereas the average relative growth inhibition of *Phy. infestans* by *Pho. eupatorii* was 50.6 ± 2.2%, the average relative inhibition of *Pho. eupatorii* by *Phy. infestans* was 4.7 ± 0.9%.

After eight weeks, the endophytes, (except for isolate 9907) visually overgrew the plates, including the regions colonized by *Phy. infestans* (Fig. 4). To substantiate this observation, we extracted DNA from some co-cultures with *Phi. fortinii* (12 co-cultures), *Pho. eupatorii* (18 co-cultures) and *Monosporascus* sp. (seven co-cultures) from both sides of the eight-week samples (Tab. S6). In total, we analyzed 37 co-cultures and their respective controls for the presence of endophyte and *Phy. infestans.* We used the marker genes *COX* and *ITS*. Because our *ITS* primers were specific for fungi, we primarily observed amplicons from the fungal endophyte *ITS* loci when both organisms were present. However, presence of *Phy. Infestans* could be determined by the presence of a *COX* amplicon. In general, we observed that the endophyte was present on both sides of the plates, whereas *Phy. infestans* was either not detected or only on the side of the plate on which it had been inoculated. Few exceptions occurred in which *Phy. infestans* was observed on the side of original inoculation of the fungal endophyte (2/37 cases). Hence, *Phy. infestans* was usually not able to colonize the side of the plate where the endophyte was growing, while the endophyte was always able to colonize *Phy. infestans’* side of the plate. In addition, the endophytes showed a greater inhibition of *Phy. infestans* than *Phy. infestans* did on the endophytes. Therefore, resource limitation (due to the size of the plates) is unlikely to fully explain the unequal growth differential between *Phy. infestans* and the endophytes during co-cultivation. Instead, we hypothesize that factors actively secreted by the endophytes may also be involved in the growth inhibition of *Phy. infestans.*

**Figure 4.**
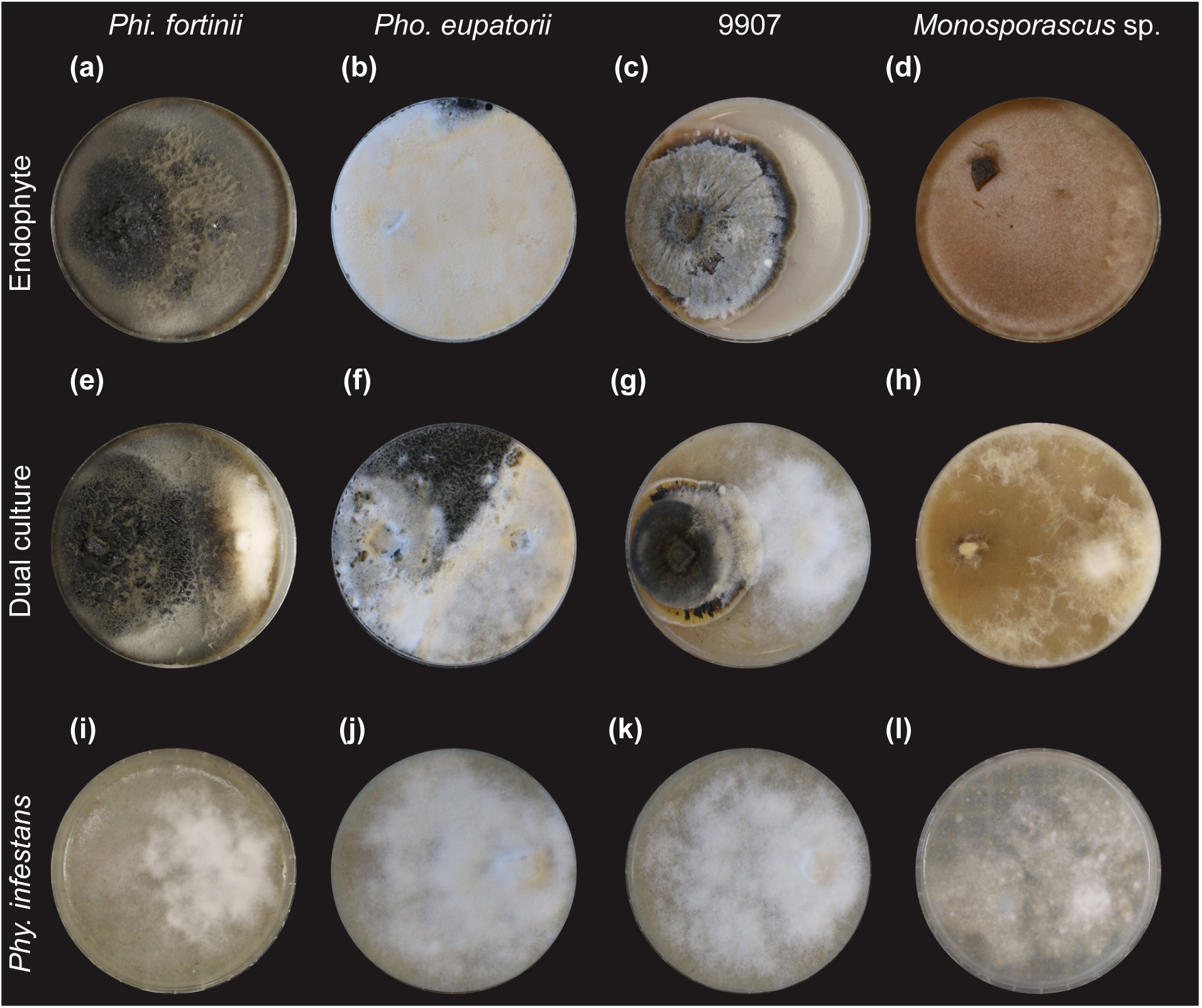
Long-term co-cultivation of fungal endophytes with *Phytophthora infestans* on plate. Examples of eight-week-old co-cultivations and their respective controls. *Phi. fortinii* with *Phy. infestans* isolate NL88069 (a, e, i), *Pho. eupatorii* with *Phy. infestans* isolate NL88069 (b, f, j), isolate 9907 with *Phy. infestans* isolate T15-2 (c, g, k) and *Monosporascus* sp. with *Phy. infestans* isolate NL10001 (d, h, l). The diameter of each plate is nine cm.

### *Phoma eupatorii* limits *Phytophthora infestans* infection success

We identified global, non-isolate-specific growth inhibition by all four endophytes in plate assays. To test whether the inhibitory potential of the endophytes holds true *in planta,* we inoculated the fungal endophytes in axenically grown *S. lycopersicum* cv. M82 seedlings. Our preliminary screening showed that *Phi. fortinii* and isolate 9907 were too virulent and killed the *S. lycopersicum* seedlings (Fig. S2a,b,d). In contrast, *S. lycopersicum* seedlings inoculated with *Pho. eupatorii* and *Monosporascus* sp. survived (Fig. S2a,c,e).

To confirm the endophytic colonization of the roots, we analyzed fungal outgrowth of surface sterilized roots and their imprints from inoculations with water, endophyte or endophyte and *Phy. infestans* (Tab. 2). Irrespective of the protocol, there was neither fungal growth from the surface sterilized mock control roots nor from their imprints. Generally, imprints of the surface sterilized endophyte inoculated roots did not show fungal growth, except for *Pho. eupatorii* inoculated roots after sterilization treatment 1 (1/16 imprints from the mono-inoculation and 5/12 imprints from the co-inoculations). This suggests that surface sterilization was successful in all other cases. *Pho. eupatorii* grew from several roots independently of the sterilization treatment, although the stronger treatments resulted in less outgrowth. Hence, these treatments may partially impact survival of endophytic mycelium. Nevertheless, these results show that *Pho. eupatrorii* is capable of colonizing *S. lycopersicum* roots. *Monosporascus* sp. also showed outgrowth from several of the plated surface sterilized roots, suggesting that, like *Pho. eupatorii, Monosporascus* sp. also grows endophytically in the roots of *S. lycopersicum. S. lycopersicum* seedlings colonized by *Pho. eupatorii* are visually smaller than mock control seedlings and seedlings mono-inoculated with *Phy. infestans*. We also observed a reduction in leaflet number (Fig. S3a,c). Since the leaflets appeared sturdier and were darker green than the controls (Fig. 5a-f), we measured chlorophyll levels via chlorophyll fluorescence. However, chlorophyll abundance did not change following any of the treatments (Fig. 5g-m). We also observed that some of the stems of the plants that had been inoculated with *Pho. eupatorii* developed a purple color (Fig. S4c). Therefore, we reasoned that the darker leaflet color may have resulted from anthocyanin accumulation. In fact, we detected a significant increase in anthocyanin content in *Pho. eupatorii* inoculated *versus* mock control plants (p=0.001 without *Phy. infestans*, p=0.04 with *Phy. infestans*, Fig. 5o). In contrast to seedlings colonized by *Pho. eupatorii*, those inoculated with *Monosporascus* sp. did not visibly differ from the mock controls (Fig. S3a,b,S4a,c). In agreement with this, anthocyanin content did not increase significantly in *Monosporascus* sp. inoculated samples compared to the mock control (p=0.08 without *Phy. infestans*), but slightly when both endophyte and pathogen were present (p=0.007), suggesting that the increase results from the presence of *Phy. infestans*.

**Figure 5.**
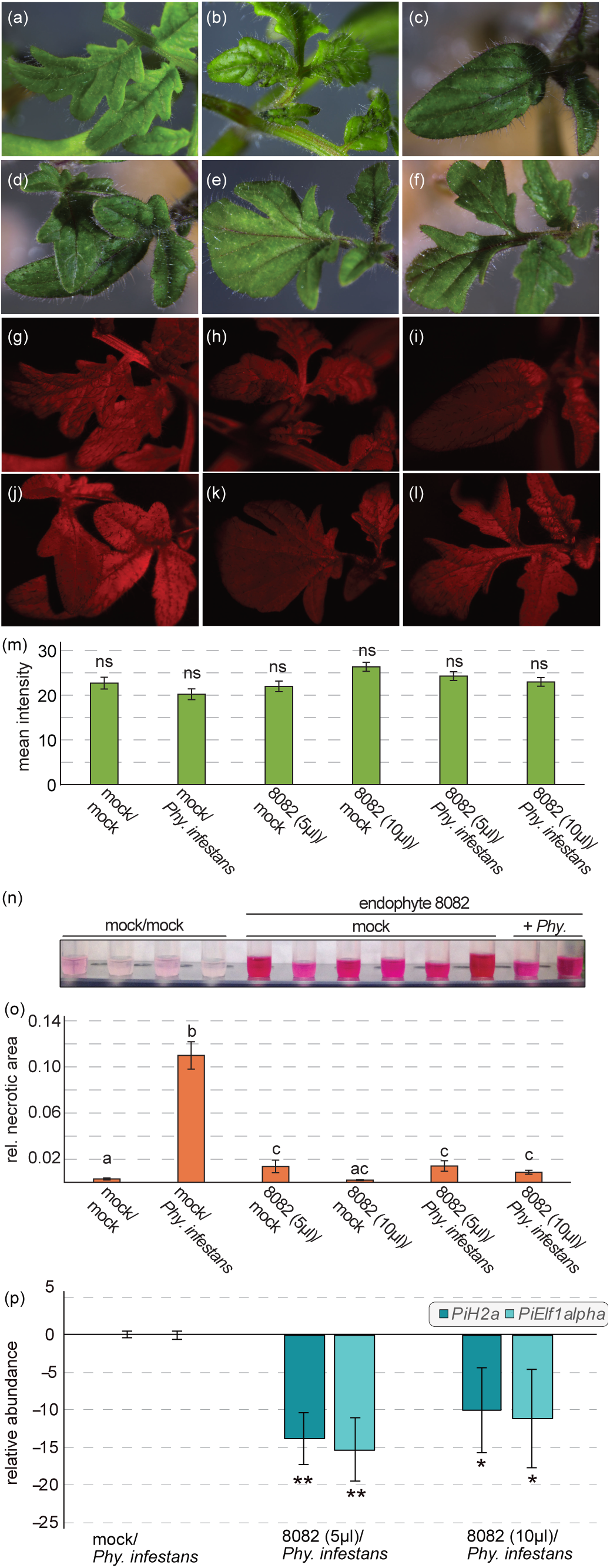
*In planta* co-inoculations of *Phoma eupatorii* isolate 8082 and *Phytophthora infestans*. *S. lycopersicum* cv. M82 seedlings were mock treated (a) or inoculated with *Phy*. *infestans* isolate D12-2 (b), 5μl of *Pho*. *eupatorii* mycelium suspension (c), 10μl of *Pho. eupatorii* mycelium suspension (d), 5μl of *Pho. eupatorii* mycelium suspension and *Phy*. *infestans* isolate D12-2 (e) and 10μl of *Pho. eupatorii* mycelium suspension and *Phy. infestans* isolate D12-2 (f). Chlorophyll fluorescence is depicted in red false coloring for all combinations (g-l) and was measured as mean fluorescence intensity using ImageJ (m). Bars give the average mean fluorescence (n_leaflets_=17-37). Error bars give the standard error (SEM); ns = not significant. Differences in anthocyanin content (n). A darker pink in the examples shown indicates a higher amount of anthocyanins in the sample. The average relative necrotic area of the leaflets was calculated for each treatment (n_leaflets_=38-156, o). Bars give the average necrotic area per treatment and error bars indicate the SEM. Significant differences between the treatments are indicated by different letters above the bars with a cutoff of p<0.05; same letter = not significant. The relative abundance of *Phy*. *infestans* isolate D12-2 was measured with a qRT-PCR of the two biomass marker genes *PiH2a* and *PiElf1 α* (p). Bars show average relative expression of the two biomass markers normalized against the three plant reference genes *SAND, TIP* and *TIF3H* and compared between the *Pho. eupatorii* –*Phy. infestans* co-inoculations and the control treatment (*Phy. infestans* only). The error bars indicate the SEM. Significant differences between relative *Phy. infestans* abundance in samples pre-inoculated with the endophyte and the control are indicated by * p<0.05 and ** p< 0.01. In all bar graphs, treatments with *Pho. eupatorii* are indicated by its isolate number 8082.

**Table 2.** Endophytic outgrowth from surface sterilized roots after inoculation with the endophyte. Roots were surfaces sterilized and an imprint of each root was prepared to test for efficiency of the treatment. The days after which the roots were surveyed is given as days post sterilization (dps). Treatment 1, 2 and 3 indicate the type of surface sterilization as described in the Material and Method section. The number of imprints and roots with fungal growth and the total number of analyzed roots is given for each sample type.

|  | *Pho. eupatorii* 8dps |  | *Monosporascus* sp. 6dps |  |
| --- | --- | --- | --- | --- |
|  | imprint | roots | imprint | roots |
| **Treatment 1** |  |  |  |  |
| mock/mock | 0/10 | 0/10 | 0/13 | 0/13 |
| endophyte/mock | 1/16 | 13/16 | 0/12 | 3/12 |
| endophyte/*Phy. infestans* | 5/12 | 10/12 | 0/12 | 3/12 |
| **Treatment 2** |  |  |  |  |
| mock/mock | 0/10 | 0/10 | 0/12 | 0/12 |
| endophyte/mock | 0/13 | 2/13 | 0/12 | 3/12 |
| endophyte/*Phy. infestans* | 0/12 | 3/12 | 0/12 | 0/12 |
| **Treatment 3** |  |  |  |  |
| mock/mock | 0/11 | 0/11 | 0/12 | 0/12 |
| endophyte/mock | 0/15 | 4/15 | 0/12 | 0/12 |
| endophyte/*Phy. infestans* | 0/12 | 2/12 | 0/8 | 2/8 |

Despite the visible effects of the colonization by *Pho. eupatorii* on the seedlings, we proceeded to investigate the effect of the endophyte on a subsequent infection with *Phy. infestans*. The relative necrotic area caused by the pathogen is significantly higher on plants inoculated only with *Phy. infestans* (in the absence of pre-inoculation by an endophyte) compared to the mock control (Fig. 5n,S4e). To confirm the pathogen infection in the mock/*Phy. infestans* samples, we used the expression of the *Phy. infestans* biomass marker genes *PiH2a* and *PiElf1α*. In agreement with the increase in necrotic area, *Phy. infestans* was present in all biological replicates mono-inoculated with the pathogen, i.e. demonstrating a successful infection.

While the relative necrotic area in seedlings that were colonized only by *Pho. eupatorii* was 4.7-fold higher compared to the mock control, this was significantly less than the relative necrotic area of seedlings infected with only *Phy. infestans* (Fig. 5n). *S. lycopersicum* seedlings co-inoculated with *Pho. eupatorii* and *Phy. infestans* resulted in a significantly reduced relative necrotic area compared to seedlings mono-inoculated with *Phy. infestans* (Fig. 5n). Importantly, the average relative necrotic area of leaflets colonized by both *Pho. eupatorii* and *Phy. infestans* did not differ from the mono-inoculations with the endophyte (Fig. 5n). Whether 5μl or 10μl mycelial suspensions of *Pho. eupatorii* was used had no effect on the outcome of the experiments. The relative necrotic area between the treatment with *Monosporascus* sp. and the mock control did not differ (Fig. S4a,c,e). This endophyte was not able to inhibit *Phy. infestans* infection nor limit development of disease symptoms *in planta* (Fig. S4b,d,e,f).

To quantify the biomass of *Phy. infestans in planta* after pre-inoculation with *Pho. eupatorii,* we performed a qRT-PCR with the two biomass marker genes *PiElf1α* and *PiH2A* (Fig. 5o). In total, we tested the three biological replicates from the 5μl *Pho. eupatorii* inoculations and two from the 10μl *Pho. eupatorii* inoculations. In three of those five replicates, we did not detect an amplicon for either *PiH2a* or *PiElf1α.* Yet, *PiH2a* and *PiElf1α* were detected in every biological replicate of the mock/*Phy. infestans* infections. In addition, three plant-specific reference genes were tested; these showed no aberrant expression in any of the samples colonized by the endophyte in which *PiH2a* and *PiElf1α* were not detected. Hence the presence of the fungal endophyte did not affect the efficiency of the qRT-PCR. Also, those samples that were pre-inoculated with *Pho. eupatorii,* but gave an amplicon of the marker genes had reduced Cq-values for both marker genes compared to the mock/*Phy. infestans* samples. This suggests that *Pho. eupatorii* reduced the infection with *Phy. infestans* isolate D12-2 in the sampled leaflets. To estimate the reduction of *Phy. infestans* biomass, we assumed that the Cq-value of those replicates with no amplicon could theoretically have been amplified in later cycles. We therefore set the Cq-values in those samples to 41; i.e. one cycle more than the original runs included. Based on this assumption, we observed a significant reduction of gene expression in both biomass marker genes in the *Pho. eupatorii* pre-treated samples compared to mono-infections of *Phy. infestans* (Fig. 5o). Therefore, *Pho. eupatorii* is capable of significantly inhibiting *Phy. infestans* infection of *S. lycopersicum* leaflets.

## Discussion

### Fungal endophytes show a broad-spectrum growth inhibition of European *Phy. Infestans* isolates

Of 12 fungi for which culture extracts were tested for inhibition of *Phy. infestans,* we identified three ascomycetes, *Pho. eupatorii,* isolate 9907 and *Monosporascus* sp., which effectively inhibited growth of the pathogen. While fungal endophytes produce a vast diversity of metabolites (Schulz *et al.*, 2002; Strobel & Strobel, 2007; Verma *et al.*, 2009; Mousa & Raizada, 2013; Brader *et al.*, 2014) and numerous have antimicrobial activity (Son *et al.*, 2008; Puopolo *et al.*, 2014; Mousa *et al.*, 2016), endophytes and their metabolites may have a narrow spectrum of specificity. To avoid narrow spectrum of pathogen inhibition, we screened these three fungal endophytes and the endophyte *Phi. fortinii* for their capacity to inhibit the growth of nine European isolates of *Phy. infestans*. In our co-culture assays, *Pho. eupatorii* and isolate 9907 had a broad-spectrum inhibition against all tested isolates, whereas *Monosporascus* sp. and *Phi. fortinii* inhibited most of the isolates. Additionally, after eight-weeks of incubation, the pathogen was not able to grow on sections of the plates, in which the endophytes grew. The consistency of the results from the culture extract experiments and the plate assays of *Pho. eupatorii* and isolate 9907 shows that their inhibition is independent of the growth medium, suggesting an environmentally robust metabolite production of their anti-*Phytophthora* substances. A robust metabolite production would be of great advantage, if these fungal endophytes are to be used as living biocontrol agents in the field.

For application in the field, two issues must be examined: i) Does infection by the endophyte damage the host in the absence of a pathogen? and ii) Does the endophyte successfully inhibit the pathogen in the host? In our study, the former is of extreme importance, because the fungal endophytes in question were not originally isolated from the Solanaceae, to which tomato belongs. Furthermore, whether an endophyte remains benign and asymptomatic is likely to be affected by a number of different circumstances and in some cases the host endophyte relationship may shift to a pathogenic outcome from an initially protective interaction (Schulz & Boyle, 2005; Junker *et al.*, 2012; Schulz *et al.*, 2015; Busby *et al.*, 2016b). Along these lines we excluded two isolates, *Phi. fortinii* and isolate 9907, for direct applications as biocontrol agents: Seedlings of *S. lycopersicum* infected with either of these two isolates quickly died after inoculation. A third isolate, *Monosporascus* sp., neither inhibited *Phy. infestans* infection nor hindered its infection progress. This may not be surprising, because *Monosporascus* sp. had the lowest inhibition potential in our co-culture assays. It should, however, be noted that the metabolite composition of fungal endophytes varies depending on their environments, i.e. *in vitro* and *in planta* (Brader *et al.*, 2014). It is therefore possible that the metabolite composition *Monosporascus* sp. produces *in planta* does not include the active anti-*Phytophthora* compound. Alternatively, the active compound may be only produced in specific stages of the infection. In the latter scenario, the infection of *Monosporascus* sp. may not have progressed far enough by the time we inoculated with *Phy. infestans.* Nevertheless, the outcome of the *in planta* co-inoculations does not exclude the possibility that the *in vitro* produced metabolites could be effective in field applications, especially since they showed a broad-spectrum reduction in *Phy. infestans* growth in our co-culture experiments. The broad-spectrum effectiveness of inhibition suggests that the metabolite composition either includes a metabolite with a conserved target in *Phy. infestans* or a mixture of anti-*Phytophthora* metabolites. Both would slow the counter-adaptation of the pathogen to the metabolites if used in field application. As a next step, the metabolite extracts with protective capabilities should be tested for their cytotoxicity *in planta*.

### *Phoma eupatorii* isolate 8082 may inhibit *Phytophthora infestans* via secreted toxic metabolite(s) and/or the induction of host defense mechanisms

*Pho. eupatorii* was the most effective fungal endophyte in our experiments, excelling both in co-culture as well as *in planta*. The presence of *Pho. eupatorii* not only reduced or inhibited the pathogen’s growth, but perhaps entirely prevented infection. Here we used root inoculations of *Pho. eupatorii* combined with leaflet inoculations of *Phy. infestans* isolate D12-2. Because *Pho. eupatorii* was applied to roots, while *Phy. infestans* was inoculated on the leaves, niche competition is an unlikely mechanism by which *Pho. eupatorii* protects the *S. lycopersicum* seedlings. Therefore, two other possible mechanisms by which the plant is defended against the pathogen include endophyte-dependent induction of defense responses or the production of anti-*Phytophthora* metabolites. The induction of plant defense responses by endophytes, such as *Pir. indica* and non-pathogenic *Fusarium oxysporum*, has been previously shown (Stein *et al.*, 2008; Aimé *et al.*, 2013). Here, we observed an elevation of anthocyanin levels in leaf tissue of *S. lycopersicum* after root colonization of *Pho. eupatorii.* Accumulation of anthocyanins is, among other factors, positively regulated by jasmonic acid (Franceschi & Grimes, 1991; Feys *et al.*, 1994; Shan *et al.*, 2009; Li *et al.*, 2006). Hence, it is possible that jasmonic acid dependent defense responses are induced upon colonization with *Pho. eupatorii*. This may be a response to *Pho. eupatorii* and elevated levels of jasmonic acid may have contributed to the inhibition of the *Phy. infestans* infection we observed. The role of jasmonic acid in defense against *Phy. infestans* is not clear: In one study, application of jasmonic acid to leaves of tomato and potato plants resulted in reduced infection of the pathogen (Cohen *et al.*, 1993). In another study, it is reported that jasmonic acid is required for the initiation of defense responses triggered by a peptide secreted by *Phy. infestans* (Halim *et al.*, 2009). Yet, potato RNA interference lines that downregulated jasmonic acid biosynthesis and signaling components, showed no alterations in the infection rates of *Phy. infestans* (Halim *et al.*, 2009). Hence, the production of anti-*Phytophthora* metabolites may be a more likely explanation for the observed reduction of *Phy. infestans* infection. A recently published example of a metabolite based endophyte-mediated pathogen protection is that of *Enterobacter* sp. This endophyte produces many different antimicrobial compounds in its host plant and these are detrimental to the host plant’s pathogen *F. graminearum* (Mousa *et al.*, 2016). In our study, each of the four fungal endophytes undoubtedly produces anti-*Phytophthora* metabolites in the crude extract tests and in the co-cultures on agar media. This makes it likely that *Pho. eupatorii* also produces such metabolites during *in planta* co-inoculations with *Phy. infestans.* A combination of these two mechanisms is, however, also possible.

### Conclusion: *Phoma eupatorii* isolate 8082 is a potential novel *Phytophthora infestans* biocontrol agent

Out of a screen of 12 fungal endophytes, we discovered four ascomycetes that inhibited the growth of *Phy. infestans* in co-culture, presumably through the secretion of secondary metabolites, particularly since their culture extracts were also active. Most importantly, two of the endophytes exhibited global inhibition of nine European *Phy. infestans* isolates, the other two showing a near-global inhibition. This indicates that a conserved target within *Phy. infestans* for a particular metabolite may be produced by these four endophytes. Alternatively, a complex metabolite mixture could be involved. In either case, the use of these fungi for biocontrol could slow the counter-adaptation of *Phy. infestans*. Hence, all four fungal endophytes can be considered good candidates for the production of such new and urgently needed compounds. Additionally, of the four fungal endophytes, *Pho. eupatorii* functioned as an effective biocontrol agent *in planta.* Therefore, *Pho. eupatorii* may not only synthesize a reservoir of highly useful antimicrobial metabolites, but could serve as a novel biocontrol agent providing an alternative to resistance gene breeding and application of agrochemicals.

## Acknowledgements

We thank Tuba Altinmakas, Melissa Mantz and Klaudia Maas-Kantel for technical support, Siegfried Draeger for isolation of the endophytes and initial identification based on morphology. We thank the TGRC Institute for providing the seeds of *S. lycopersicum* cv. M82 and Francine Govers (Wageningen University) for the *Phy. infestans* isolates NL10001, NL88069, NL90128, IPO-C, IPO428-2, 3928A, D12-2, T15-2 and T20-2. We thank Dr. Bärbel Schöber-Butin for providing the German isolate *Phy. infestans* D2. This work was supported by the Deutsche Forschungsgemeinschaft (Ro 2491/5-2, Ro 2491/6-1, Research Training Group GRK1525).

## Author contribution

SdV, BS and LER wrote the manuscript. SdV, JKvD, AS and SG performed the experimental work and data analyses. BS provided the fungal isolates and the metabolite screening. All authors read and approved the manuscript.

## References

Adomas A, Eklund M, Johansson M, Asiegbu FO. 2006. Identification and analysis of differentially expressed cDNAs during nonself-competitive interaction between *Phlebiopsis gigantea* and *Heterobasidion parviporum*. FEMS Microbiology Ecology 57: 26–39.

Aimé S, Alabouvette C, Steinerg C, Olivain C. 2013, The endophytic strain *Fusarium oxysporum* Fo47: A good candidate for priming the defense responses in tomato roots. Molecular Plant-Microbe Interactions 26: 918–926.

Alabouvette C, Olivain C, Migheli Q, Steinberg C. 2009, Microbiological control of soil-borne phytopathogenic fungi with special emphasis on wilt-inducing *Fusarium oxysporum*.New Phytologist 184: 529–544.

Altschul SF, Gish W, Miller W, Myers EW, Lipman DJ. 1990. Basic local alignment search tool. Journal of Molecular Biology 215: 403–410.

Annesi T, Curcio G, D’Amico L, Motta E. 2005, Biological control of *Heterobasidion annosum* on *Pinus pinea* by *Phlebiopsis gigantea*. Forest Pathology 35: 127–134.

Aveskamp MM, de Gruyter J, Woudenberg JHC, Verkley GJM, Crous PW. 2010. Highlights of the Didymellaceae: A polyphasic approach to characterise *Phoma* and related pleosporalean genera. Studies in Mycology 65: 1–60.

Bahramisharif A, Lamprecht SC, Calitz F, McLeod A. 2013. Suppression of *Pythium* and *Phytophthora* damping-off of rooibos by compost and a combination of compost and non-pathogenic *Pythium*-taxa. Plant disease 97: 1605–1610.

Benjamini Y, Hochberg Y. 1995. Controlling the false discovery rate: a practical and powerful approach to multiple testing. Journal of the Royal Statistical Society Series B 57:289–300.

Brader G, Compant S, Mitter B, Trognitz F, Sessitsch A. 2014. Metabolic potential of endophytic bacteria. Current Opinion in Biotechnology 27: 30–37.

Bodenhausen N, Horton MW, Bergelson J. 2013. Bacterial communities associated with leaves and the roots of *Arabidopsis thaliana*. Plos One 8: e56329.

Bulgarelli D, Rott M, Schlaeppi K, Ver Loren van Themaat E, Ahmadinejad N, Assenza F,Rauf P, Huettel B, Reinhardt R, Schmelzer E, Peplies J, et al. 2012. Revealing structure and assembly cues for *Arabidopsis* root-inhabiting bacterial microbiota.Nature 488: 91–95.

Bulgarelli D, Garrido-Oter R, Münch PC, Weiman A, Dröge J, Pan Y, McHardy AC, Schulze-Lefert P. 2015. Structure and function of the bacterial root microbiota in wild and domesticated barley. Cell Host and Microbe 17: 392–403.

Busby PE, Peay KG, Newcombe G. 2016a. Common foliar fungi of *Populus trichocarpa* modify *Melampsora* rust disease severity. New Phytologist 209: 1681–1692.

Busby PE, Ridout M, Newcombe G. 2016b. Fungal endophytes: modifiers of plant disease. Plant Molecular Biology 90: 645–655.

Cao L, Qiu Z, You J, Tan H, Zhou S. 2004. Isolation and characterization of endophytic *Streptomyces* strains from surface-sterilized tomato (*Lycopersicon esculentum*) roots. Letter in Applied Microbiology 39: 425–430.

Caten CE, Jinks JL. 1968. Spontanous variability of single isolates of *Phytophthora infestans.* I. Culture variation. Canadian Journal of Botany 46: 329–348.

Childers R, Danies G, Myers K, Fei Z, Small IM, Fry WE. 2015. Acquired resistance to mefenoxam in sensitive isolates of *Phytophthora infestans*. Phytopathology 105: 342–349.

Collado J, Gonzalez A, Platas G, Stchigel AM, Guarro J, Pelaez F. 2002. *Monosporascus ibericus* sp. nov. an endophytic ascomycete from plants on saline soils, with observations on the position of the genus based on sequence analysis of the 18S rDNA. Mycological Research 106: 118–127.

Cohen Y, Gisi U, Niderman T. 1993. Local and systemic protection against *Phytophthora infestans* induced in potato and tomato plants by jasmonic acid and jasmonic methyl ester. Phytopathology 83: 1054–1062.

Coleman-Derr D, Desgarennes D, Fonseca-Garcia C, Gross S, Clingenpeel S, Woyke T, North G, Visel A, Partida-Martinez LP, Tringe SG. 2016. Plant compartment and biogeography affect microbiome composition in cultivated and native *Agave* species. New Phytologist 209: 798–811.

Crous PW, Gams W, Stalpers JA, Robert V, Stegehuis G. 2004. MycoBank: an online initiative to launch mycology into the 21st century. Studies in Mycology 50: 19–22.

Edwards J, Johnson C, Santos-Medellín C, Lurie E, Podishetty NK, Bhatnagar S, Eisen JA, Sundaresan V. 2015. Structure, variation, and assembly of the root-associated microbiomes of rice.Proceedings of the National Academy of Sciences of the United States of America 112: E911–E920.

Feys BJ, Benedetti CE, Penfold CN, Turner JG. 1994. *Arabidopsis* mutants selected for resistance to the phytotoxin coronatine are male sterile, insensitive to methyl jasmonate and resistant to a bacterial pathogen. The Plant Cell 6: 751–759.

Franceschi VR, Grimes HD. 1991. Induction of soybean vegetative storage proteins and anthocyanins by low-level atmospheric methyl jasmonate.Proceedings of the National Academy of Sciences of the United States of America 88: 6745–6749.

Grünig CR, Queloz V, Sieber TN, Holdenrieder O. 2008.Dark septate endophytes (DSE) of the *Phialocephala fortinii* s.l. – *Acephala applanata* species complex in tree roots: classification, population biology, and ecology. Botany 86: 1355–1369.

Grünwald NJ, Sturbaum AK, Montes GR, Serrano EG, Lozoya-Saldaña H, Fry WE. 2006. Selection for fungicide resistance within a growing season in field populations of *Phytophthora infestans* at the center of origin. Phytopathology 96: 1397–1403.

Halim VA, Altmann S, Ellinger D, Eschen-Lippold L, Miersch O. 2009. PAMP-induced defense responses in potato require both salicylic acid and jasmonic acid. The Plant Journal 57: 230–242.

Hiruma K, Gerlach N, Sacristán S, Nakano RT, Hacquard S, Kracher B, Neumann U, Ramírez D, Bucher M, O’Connell RJ, et al. 2016.Root endophyte *Colletrichum tofieldiae* confers plant fitness benefits that are phosphate status dependent. Cell 165: 464–474.

Höller U, Wright AD, Matthée GF, König GM, Draeger S, Aust H-J, Schulz B. 2000. Fungi from marine sponges: diversity, biological activity and secondary metabolites. Mycological Research 104: 1354 - 1365.

Hudspeth DSS, Nadler SA, Hudspeth MES. 2000. A COX2 molecular phylogeny of the Peronosporomycetes. Mycologia 92: 674–684.

Junker C, Draeger S, Schulz B. 2012. A fine line – endophytes or pathogens in *Arabidopsis thaliana*. Fungal Ecology 5: 657 - 662.

Kim H-Y, Choi GJ, Lee HB, Lee S-W, Lim HK, Jang KS, Son SW, Lee SO, Cho KY, Sung ND, et al. 2007. Some fungal endophytes from vegetable crops and their anti-oomycete activities against tomato late blight. Letters in Applied Microbiology.44: 332–337.

Kruskal WH, Wallis WA. 1952. Use of ranks in one-criterion variance analysis. Journal of the American Statistical Association 47: 583–621.

Lahlali R, Hijri M. 2010. Screening, identification and evaluation of potential biocontrol fungal endophytes against *Rhizoctonia solani* AG3 on potato plants.FEMS Microbiology Letters. 311: 152–159.

Le Cocq K, Gurr SJ, Hirsch PR, Mauchline TH. 2016. Exploiting of endophytes for sustainable agriculture intensification. Molecular Plant Pathology doi:10.1111/mpp.12483.

Li J, Brader G, Kariola T, Palva ET. 2006. WRKY70 modulates the selection of signaling pathways in plant defense. The Plant Journal 46: 477–491.

Lindoo SJ, Caldwell MM. 1978.Ultraviolet-B radiation-induced inhibition of leaf expansion and promotion of anthocyanin production. Plant Physiology 61: 178–282.

Lundberg DS, Lebeis SL, Paredes SH, Yourstone S, Gehring J, Malfatti S, Tremblay J, Engelbrekston A, Kunin V, Glavina del Rio T, et al. 2012. Defining the core *Arabidopsis thaliana* root microbiome. Nature 488: 86–90.

Mann HB, Whitney DR. 1947. On a test of whether one of two random variables is stochastically larger than the other. Annals of Mathematical Statistics 1: 50–60.

Martínez-Medina A, Fernandez I, Lok GB, Pozo MJ, Pieterse CMJ, Van Wees SCM. 2017. Shifting from priming of salicyliuc acid-to jasmonate acid-regulated defences by *Trichoderma* protects tomato against the root knot nematode *Meloidogyne incognita*. New Phytologist 213: 1363–1377.

Miles LA, Lopera CA, González S, Cepero de García MC, Franco AE, Restrepo S. 2012. Exploring the biocontrol potential of fungal endophytes from an Andean Colombian paramo ecosystem. BioControl 57: 697–710.

Mousa WK, Raizada MN. 2013. The diversity of anti-microbial secondary metabolites produced by fungal endophytes: an interdisciplinary approach. Frontiers in Microbiology 4: 65.

Mousa WK, Shearer C, Limay-Rios V, Ettinger CL, Eisen JA, Raizada MN. 2016. Root-hair endophyte stacking in finger millet creates a physiochemical barrier to trap the fungal pathogen *Fusarium graminearum*. Nature Microbiology 1: 16167.

Murashige T, Skoog F. 1962. A revised medium for rapid growth and bio assays wity tobacco tissue cultures. Physologia Plantarum 15: 473–497.

Nowicki M, Foolad MR, Nowakowska M, Kozik EU. 2012. Potato and tomato late blight caused by *Phytophthora infestans*: an overview of pathology and resistance breeding. Plant disease 96: 4–17.

Panke-Buisse K, Poole AC, Goodrich JK, Ley RE, Kao-Kniffin J. 2015. Selection on soil microbiomes reveals reproducible impacts on plant function. The ISME Journal 9: 980–989.

Pel MA, Foster SJ Park T-H, Rietman H, van Arkel G, Jones JDG, van Eck HJ, Jacobsen E, Visser RGF, van der Vossen E. 2009. Mapping and cloning of late blight resistance genes from *Solanum venturii* using an interspecific candidate gene approach. Molecular Plant-Microbe Interactions 22: 601–615.

Peters S, Aust H-J, Draeger S, Schulz B. 1998. Interactions in dual cultures of endophytic fungi with host and nonhost plant calli. Mycologia 90: 360–367.

Pfaffl MW. 2001. A new mathematical model for relative quantification in real-time RT-PCR. Nucleic Acids Research 29: e45.

Ploch S, Rose LE, Bass D, Bonkowski M. 2016. High diversity revealed in leaf-associated protists (Rhizaria: Cercozoa) of Brassicaceae. Journal of Eukaryotic Microbiology 63: 635–641.

Puopolo G, Cimmino A, Palmieri MC, Giovannini O, Evidente A, Pertot I. 2014. *Lysobacter capsici* AZ78 produces cyclo(L-Pro-L-Tyr), a 2,5-diketopiperazine with toxic activity against sporangia of *Phytophthora infestans* and *Plasmopara viticola*. Journal of Applied Microbiology 117: 1168–1180.

Rolli E, Marasco R, Vigani G, Ettoumi B, Mapelli F, Deangelis ML, Gandolfi C, Casati E, Previtali F, Gerbino R. 2015. Improved plant resistance to drought is promoted by the root-associated microbiome as a water stress-dependent trait. Environmental Microbiology 17: 316–331.

Schlaeppi K, Dombrowski N, Oter RG, Ver Loren van Themaat E, Schulze-Lefert P. 2013. Quantitative divergence of the bacterial root microbiota in *Arabidopsis thaliana* relatives. Proceedings of the National Academy of Sciences of the United States of America 111: 585–592.

Schindelin J, Rueden CT, Hiner MC, Eliceiri KW. 2015. The ImageJ ecosystem: An open platform for biomedical image analysis. Molecular Reproduction and Development 82: 518–529.

Schulz B, Boyle C, Draeger S, Römmert A-K, Krohn K. 2002. Endophytic fungi: a source of novel biologically active secondary metabolites. Mycological Research 106: 996–1004.

Schulz B, Boyle C. 2005. The endophytic continuum. Mycological research 109: 661–686.

Schulz B. 2006. Mutualistic interactions with fungal root endophytes. In: Schulz BJE, Boyle CJC, Sieber TN, eds. Microbial Root Endophytes. Berlin, Germany: Springer Verlag, 261–279.

Schulz B, Krohn K, Meier K, Draeger S. 2011. Isolation of endophytic fungi for the production of biologically active secondary metabolites. In: Pirttilä AM, Sovari S, eds. Prospects and Applications for Plant-Associated Microbes. Turku, Finland: BioBien Innovations, 88–95.

Schulz B, Haas S, Junker C, Andrée N, Schobert M. 2015.Fungal endophytes are involved in multiple balanced antagonisms. Current Science 109: 39–45.

Shan X, Zhang Y, Peng W, Wang Z, Xie D. 2009. Molecular mechanism for jasmonate-induction of anthocyanin accumulation in *Arabidopsis*. Journal of Experimental Botany 60: 3849–3860.

Shapiro SS, Wilk MB. 1965. An analysis of variance test for normality (complete samples). Biometrika 52: 591–611.

Shoresh M, Harman GE, Mastouri F. 2010.Induced systemic resistance and plant responses to fungal biocontrol agents. Annual Review of Phytopathology 48: 21–43.

Son SW, Kim HY, Choi GJ, Lim HK, Jang KS, Lee SO, Lee S, Sung ND, Kim J-C. 2008. Bikaverin and fusaric acid from *Fusarium oxysporum* show antioomycete activity against *Phytophthora infestans*. Journal of Applied Microbiology 104: 692–698.

Song J, Bardeen JM, Naess SK, Raasch JA, Wielgus SM, Haberlach GT, Liu J, Kuang H, Austin-Phillips S, Buell CR, et al. 2003. Gene *RB* from *Solanum bulbocastanum* confers broad spectrum resistance to potato late blight. Proceedings of the National Academy of Sciences of the United States of America 100: 9128–9133.

Stein E, Molitor A, Kogel K-H, Waller F 2008. Systemic resistance in *Arabidopsis* conferred by the mycorrhizal fungus *Piriformospora indica* requires jasmonic acid signaling and the cytoplasmic function of NPR1. Plant and Cell Physiology 49: 1747–1751.

Strobel SA, Strobel GA. 2007. Plant endophytes as a platform for discovery-based undergraduate science education. Nature Chemical Biology 3: 356–359.

Sturz AV, Christie BR, Matheson BG, Arsenault WJ, Buchanan NA. 1999. Endophytic bacterial communities in the periderm of potato tubers and their potential to improve resistance to soil-borne plant pathogens. Plant Pathology 48: 360–369.

Suryanarayanan TS, Govinda Rajul MB, Vidal S. 2016.Biological control through fungal endophytes: gaps in knowledge hindering success. Current Biotechnology 5: doi 10.2174/2211550105666160504130322.

Tamura K, Peterson D, Peterson N, Stecher G, Nei M, Kumar S. 2011.MEGA5: molecular evolutionary genetics analysis using maximum likelihood, evolutionary distance and maximum parsimony methods. Molecular Biology and Evolution 28: 2731–2739.

Tellenbach C, Sieber TN. 2012. Do colonization by dark septate endophytes and elevated temperature affect pathogenicity of oomycetes? FEMS Microbiology Ecology 82: 157–168.

Terhonen E, Sipari N, Asiegbu FO. 2016. Inhibition of phytopathogens by fungal root endophytes of Norway spruce. Biological Control 99: 53–63.

Tukey J. 1949. Comparing individual means in the analysis of variance. Biometrics 5: 99–114.

Verma VC, Kharwar RN, Strobel GA. 2009. Chemical and functional diversity of natural products from plant associated endophytic fungi. Natural Product Communications 4: 1511–1532.

Vleeshouwers VG, Raffaele S, Vossen JH, Champouret N, Oliva R, Segretin ME, Rietman H, Cano LM, Lokossou A, Kessel G, et al. 2011. Understanding and exploiting late blight resistance in the age of effectors.Annual Reviews of Phytopathology 49: 507–531.

van der Vossen E, Sikkema A, te Lintel Hekkert B, Gros J, Stevens P, Muskens M, Wouters D, Pereira A, Stiekema W, Allefs S. 2003. An ancient *R* gene from the wild potato species *Solanum bulbocastanum* confers broad-spectrum resistance to *Phytophthora infestans* in cultivated potato and tomato. The Plant Journal 36: 867–882.

de Vries S, Kloesges T, Rose LE. 2015. Evolutionarily dynamic, but robust, targeting of resistance genes by the miR482/2118 gene family in the Solanaceae. Genome Biology and Evolution 7: 3307–21.

de Vries S, von Dahlen JK, Uhlmann C, Schnake A, Kloesges T, Rose LE. 2017. Signatures of selection and host-adapted gene expression of the *Phytophthora infestans* RNA silencing suppressor PSR2. Molecular Plant Pathology 18: 110–124.

White TJ, Bruns T, Lee S, Taylor JW. 1990. Amplification and direct sequencing of fungal ribosomal RNA genes for phylogenetics. In: Innis MA, Gelfand DH, Sninsky JJ, White TJ, eds.PCR Protocols: A Guide to Methods and Applications. NewYork, USA: AcademicPress, Inc., 315–322.

Zhang Y, Schoch CL, Fournier J, Crous PW, de Gruyter J, Woudenberg JHC, Hirayama K, Tanaka K, Pointing SB, Spatafora JW, et al. 2009. Multi-locus phylogeny of Pleosporales: a taxonomic, ecological and evolutionary re-evaluation. Studies in Mycology 64: 85–102.

Zhang C, Liu L, Zheng Z, Sun Y, Zhou L, Yang Y, Cheng F, Zhang Z, Wang X, Huang S, et al. 2013. Fine mapping of the *Ph-3* gene conferring resistance to late blight (*Phytophthora infestans*) in tomato. Theoretical and Applied Genetics 126: 2643–2653.

